# Quantifying the impacts of urban human-wildlife conflicts – how can we solve the urban gull problem in St Andrews?

**DOI:** 10.1101/2020.09.01.276899

**Authors:** Grania P. Smith

## Abstract

Interactions between urban wildlife and people are increasing globally. Some of these interactions can be negative and lead to human-wildlife conflicts. In St Andrews, Scotland, residents and business owners have come into conflict with herring gulls (*Larus argentatus*) and lesser black-backed gulls (*Larus fuscus*) that nest and forage in the town. This study quantified the number, species and distribution of nesting gulls; the vulnerability of different sources of rubbish to attack; and the likelihood of negative human-gull interactions related to food. Surveys were conducted in St Andrews during the 2016 breeding season (May-July). Nesting gull density and distribution were estimated during weekly street surveys of buildings; vantage surveys were conducted for some buildings and a correction factor estimating a minimum number of nesting gulls was produced. 110 nesting gull pairs were estimated and these occupied ~10% of buildings. The vulnerability of waste sources to attack was monitored during transects recording whether or not rubbish sources were attacked. Black bin bags had the highest probability of being attacked, but placing these in secured hessian bags prevented this. The frequency of negative human-gull interactions involving food at street-level was determined during 10-minute timed watches at various locations. Incidences of gulls taking food were rare; only eight were seen in 30 hours of watches. Altering human behaviour (for example, disposing of waste securely) will mitigate potential issues with urban wildlife. Findings from this study will enable effective management of human-gull conflicts in St Andrews and have potential applications in other urban communities.

## 1. Introduction

Human-wildlife conflicts are diverse, ranging across a multitude of landscapes from rural issues with livestock loss, to urban problems involving property damage (Rock, 2005; Michalski et al., 2006; Gusset et al., 2009; Soulsbury and White, 2016). These conflicts are defined as negative interactions between humans and wildlife, where an action by either has a detrimental impact on the other (Conover, 2001). Managing conflicts that arise between people and wildlife can be difficult, but where possible, quantifying these at a local scale can aid development of effective management to mitigate problems (Conover, 2001; Young et al., 2016).

As urban environments continue to grow, the potential for interactions between people and wildlife has increased, and subsequently the nature of urban human-wildlife conflicts have expanded (Soulsbury and White, 2016). These can include collisions between vehicles and wildlife, damage to property, spread of disease, fouling and noise, and harassment or attacks on people (Rock, 2005; Abay et al., 2011; Found and Boyce, 2011; Magle et al., 2012). Nevertheless, despite the often prominent media attention associated with these events, urban ecology is still a relatively new, albeit rapidly growing field, and urban human-wildlife conflicts remain relatively under studied (Adams, 2005; Magle et al., 2012; Soulsbury and White, 2016). It is essential that urban ecologists monitor wildlife, and work alongside others, including local councils, economists, and the public health sector to make ethical, efficient, and effective management decisions based on quantified results (Soulsbury and White, 2016). In an increasingly urbanised world finding ways to reduce conflicts between people and wildlife sharing these environments is fundamental.

Avian species appear to have adapted well to urban environments; this is probably because of their varied and flexible foraging abilities (Campbell, 2007). Gulls are opportunistic foragers and have colonised a number of coastal urban areas, and increasingly towns further inland (Rock et al., 2016). A coastal-urban interface can be a particularly favourable habitat for gulls, as it provides three sources of food – items from the marine and terrestrial systems, and from anthropogenic sources (Washburn et al., 2013). Furthermore, street lights in urban environments have allowed for night time feeding on anthropogenic sources (Rock and Vaughan, 2013), although a complete shift to feeding on anthropogenic food has not been seen and birds still exploit natural food sources (Coulson and Coulson, 2008). This is coupled with other potential benefits provided by urban environments including: refuges from natural predators and easily accessible nesting sites (Monaghan and Coulson, 1977; Raven and Coulson, 1997; Nager and O’Hanlon, 2016; Rock et al., 2016).

Both herring gulls (*Larus argentatus*) and lesser black-backed gulls (*Larus fuscus*) are seen in St Andrews town. These species have protected status in the UK, which makes lethal management (except in the interest of public health) illegal (JNCC, 2009). Herring gulls are red-listed on the UK Birds of Conservation Concern list (Eaton et al., 2015); in the last few decades they have been decreasing – particularly in their natural coastal habitats. Herring gull numbers declined by 57% between 1969-2002 across the British Isles, including the area around St Andrews (Nager and O’Hanlon, 2016). In contrast, numbers of lesser black-backed gulls in the UK have increased over the last few decades, however they are amber-listed (Eaton et al., 2015). This is because of the relative importance of the UK as a site for breeding populations; it is home to almost 40% of the world population (JNCC, 2016b). However, because of the reported extent of human-gull conflicts, conservation concern for these species can often be a contentious issue when trying to manage conflicts.

Conflicts caused by gulls in urban environments include damage to property, noise, bin-raiding, and harassment of people for food (Vermeer et al., 1988; Belant and Dolbeer, 1993; Rock, 2005). Reducing potential benefits provided by urban environments to gulls may make them less favourable, resulting in decreases in gull numbers and reducing potential for human-gull conflicts. Various management methods exist to deter nesting in urban environments. These include the use of nets and wires on buildings (Blokpoel and Tessier, 1984; Belant and Ickes, 1996), and scaring devices including mylar flags, alarm calls, and models of avian predators (Belant, 1997; Belant and Ickes, 1997; Soldatini et al., 2008). Many of these devices have been deemed ineffective for various reasons (Rock, 2012) including animals in urban environments becoming habituated to novel scarers quickly (Lowry et al., 2013). However, moving devices and altering intervals between alarm calls can reduce the speed at which animals become habituated (Bomford and O’Brien, 1990; Belant, 1997). Falconry has also been used effectively to reduce abundance of some “nuisance” birds (Atkins et al., 2017), however success with urban gulls has reportedly been limited as they apparently return to sites quickly after a falconry has stopped (Erickson et al., 1990). Furthermore, lesser black-backed gulls have been seen to aggressively attack falcons (Rock, 2005). Falconry was used in St Andrews in 2012, but ceased partially due to ineffectiveness and cost (L Adam, 2016, personal communication, 16 June). The most effective and appropriate management methods for particular locations can only be determined if the nature and extent of specific problems are known.

This study aimed to quantify three key areas of the urban “gull problem” in St Andrews to inform future management. a) The density, distribution and success of nesting gulls. b) The probability of different types of waste sources being used by foraging gulls. c) The probability of negative human-gull interactions at street-level involving food, where these occurred and what determined them. After an extensive search of the literature it was found that no studies considering this combination of different human-gull conflicts in one area had previously been conducted. As it is likely that these impacts are interconnected, it is essential that they are all considered when developing management strategies.

The first aim of this project was to ***quantify the density, distribution and success of nesting gulls in St Andrews***. Herring gulls have been shown to move significant amounts of nesting materials on to roofs which can block drains and cause damage to property and nesting gulls can become violent towards people in the proximity of their nests (Vermeer et al., 1988; Rock, 2005). No comprehensive study of roof-nesting gulls in St Andrews has previously been conducted. Studies at other locations and as part of larger surveys (such as Seabird 2000 (Mitchell et al., 2004)) have tended to rely on volunteers monitoring their roofs for gulls, meaning that where buildings were unoccupied data were not often collected (Raven and Coulson, 1997; Calladine et al., 2006). In this study, street and vantage point surveys to estimate the numbers of nesting pairs of gulls in the survey area were conducted by one surveyor. This avoided potential biases with unoccupied buildings being excluded or unreliable surveyors; it was also cost-effective compared to other survey methods such as using cherry-pickers to monitor roofs. Having estimates of nesting gulls on buildings is important for management, for example, if gulls are only present on a minority of buildings, then management at these locations will simply shift the problem to nearby roofs.

Street-level surveys were conducted weekly between May-July to coincide with the main period for gulls nesting and the presence of chicks (Perrins, 1970). Estimating gull nesting numbers can be challenging, as 100% coverage of roof space is not always possible, particularly in St Andrews where some buildings have almost no visibility because of their height and architecture. Conducting vantage point surveys of a sample of buildings allowed quantification of any limitations of the ground survey; a correction factor was then calculated and applied to street survey data to provide an estimate of nests. Furthermore, the presence of standing gulls on buildings was recorded throughout street surveys. Standing gulls may act as biological indicators of the presence of nests; they are conspicuous and therefore might provide an easy and efficient way to estimate nesting gulls from street-level in future studies.

The second aim was to ***determine the probability of attack for different waste sources in St Andrews***. Having an understanding of which waste sources are vulnerable to attack should aid effective management to remove anthropogenic food sources available to gulls from the town centre. Past literature has identified gull usage of anthropogenic food sources in general, but has not considered probability of attack for different waste sources within a town (Belant et al., 1998; Rock 2005). It is also likely that results from this part of the study will be applicable in other urban environments where similar waste sources may be available.

In St Andrews the local council have introduced “gull-proof’ hessian bags in which individuals can place black bin bags. These gull-proof bags can then be secured using ties and a Velcro flap. When used correctly these should prevent gulls or other foraging animals from accessing bin bags and spreading waste. Survey effort was focused on transects of the main two streets of St Andrews (Market St. and South St.) as these were areas that had the most cafés and shops so availability of anthropogenic food sources were at their highest. Permanent bins were included in transect surveys as these provided a constantly accessible food source to gulls. The local Clean and Green team were not aware of the locations of all permanent bins, and a map showing the locations of these in the town centre was produced to show whether there was any deficiency in their provision.

The third aim was to ***quantify the frequency and probability of negative human-gull interactions involving food, including food theft***. Gull species have been shown to exhibit plastic feeding behaviours, adapting to given scenarios as part of a successful foraging strategy; clearly demonstrating their cognitive abilities (Morand-Ferron et al., 2007). However, no existing literature has attempted to calculate a frequency of food theft from people by gulls. Surveys were conducted at locations where people were likely to consume food, and other negative interactions seen between gulls and people were also recorded, including gulls begging for food. This meant that the probability of a negative human-gull interaction occurring and variables that may impact this could be analysed. Whether food is stolen at a high rate, or whether negative media attention surrounding gull species has obscured public perceptions about these impacts, as has been seen with other human-wildlife conflicts, is unknown (Dickman, 2010).

## 2. Methods

Surveys were conducted between late May and July 2016 over a 10-week period; they involved one surveyor collecting data in St Andrews, on the East Coast of Scotland. Methods to address the three key aims of this study are outlined below. All statistics were performed using the statistical programme R (R Core Team, 2016), model fits were considered from diagnostic plots and model assumptions were met, unless otherwise stated. Model selection was undertaken using Akaike information criterion (AIC) and R^2^ values.

### a) Abundance and distribution of nesting gulls

#### i) Practical Methods

Street surveys of 1070 buildings were repeated weekly over 9 weeks; all surveys were conducted from 6am to 1pm between May and July 2016. Nesting birds were recorded on a building-by-building basis across St Andrews from The Scores in the North, to Lamond Drive in the South; City Road/ Largo Road acted as a western boundary, whilst Abbey Street/ the A197 acted as an eastern boundary (Appendix, Figure A). Nesting birds were identified using binoculars from ground level; the proportion of roof space visible was recorded (e.g. perfect visibility = 100%). The species of gull (lesser black-backed or herring gull) was recorded. High vantage points were used (The University Admission’s Building and Eden Court (The Scores), St Salvator’s Chapel tower and the University Main Library (North Street), the University Buchanan Building (Market St.), St Leonards School (The Pends), Holy Trinity Church tower (South St.), St Rule’s tower (Cathedral grounds) and Cammo Lodge (a private residence on City Road) to produce a sample of 90 buildings with better levels of visibility. This allowed complete counts of nests for some buildings where perfect visibility was possible from these vantage points. Numbers of nests seen from these vantage points were recorded and compared to those estimated from the standard street-level surveys. Vantage point surveys were conducted once for each vantage point, because of access availability. These surveys were conducted during the first two weeks of June, as this was expected to be the high point of the breeding season, by which time all nests should be *in situ*.

Chicks that became visible only as the season progressed (i.e. hidden nests often produced visible chicks) were also recorded. The number of standing gulls on each building was recorded to determine if this correlated with the number of nests. Coordinates for each building surveyed were recorded using the coordinate function on Google Earth (Google Earth, 2016).

#### ii) Analysis

The correction factor to adjust street survey counts to account for missed nests was obtained using the ground level data and data collected from the stratified samples at higher vantage points. This factor was calculated by dividing the actual number of nests seen on buildings in stratified vantage surveys, by the maximum number of nests found on a building during the 9-week ground surveys:

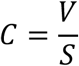

where, C = correction factor; V = nests from vantage points; and S = maximum number of nests found on corresponding buildings for street surveys.

This correction factor was then applied to all buildings that were only surveyed from ground level to produce a minimum estimate of nest numbers in St Andrews. A map of the survey area in St Andrews that could be imported into R was produced using a polygon of the survey area drawn on Google Earth (Google Earth, 2016) converted in to a shape file in the ArcGIS programme (ESRI, 2014). Data was used to produce a map of nests and chicks by overlaying building coordinates and marked nest sites on a grid format on the shape file in the statistical programme R (R Core Team, 2016); a final map incorporating a correction for the number of nests that were unrecorded from street-level surveys was produced. This included predicted missed nests, proportionally allocated to the map according to density of nests on each street, with nests placed on randomly selected buildings within those streets. The proportions of nesting lesser black-backed gulls and herring gulls was also calculated. Spatial models and Generalised Linear Models (GLMs) with a binomial error distribution were used to analyse differences in density and distribution of these species, and whether standing gulls on buildings were good indicators of nests and chicks.

### b) Factors determining whether rubbish sources are attacked by gulls

#### i) Practical Methods

After pilot studies across St Andrews, data collection was focused on South Street and Market Street because these were the areas where most restaurants and cafés in the town were present and where various vulnerable litter sources were seen. Walking transects of Market and South Street were made throughout the study at different times of day (before 07:30, afternoon (12:00-17:00) and evening (after 19:30) across all days of the week (a total of at least 21 transects per street). Each transect took approximately 30 minutes, depending on the amount of rubbish. The location, time of day, and rubbish type (gull-proof bag (tied/untied), black bin bag, permanent bin) were recorded. The locations of permanent street bins were also recorded separately to produce a map of town centre bin locations. The frequency that rubbish bags were attacked and the effect that anti-gull bags have on this frequency was calculated by recording the number of anti-gull bags and whether they were attacked or not. Attacked bin bags were checked for damage consistent with gull attacks – such as puncture marks resembling those gull bills might make. Data were also collected on fewer occasions from additional streets within and outwith the town centre (North St; Hope St; Largo Road; and Lamond Drive) to see whether litter availability was also an issue beyond the two main streets. These surveys were conducted once on a landfill bin day and once on a different day of the week.

#### ii) Analysis

A GLM with a binomial error distribution was used to determine whether different types of rubbish had different probabilities of being attacked, and results are illustrated with a graph comparing the proportion of attacks on each rubbish type. Tukey’s Post Hoc Honest Significant Differences (HSD) test and t-tests were used to see whether differences in probability of attack were significant. A map showing the location of permanent bins in the town centre was also produced to show whether there was a deficiency in available bins (Appendix, Figure B).

### c) The frequency and probability of negative human-gull interactions involving food

#### i) Practical Methods

Incidences of negative human-gull interactions including attacks on people for food were recorded during timed watches in order to determine rates for these conflicts. Timed watches of ten minutes (divided into five, two minute periods) were conducted at selected points on Market St., North St., South St., The Scores, the Cathedral grounds, and the Harbour (See Appendix, Table A for location coordinates). In total 30 hrs of watches were conducted; these were focused on Market St. and South St., as these were areas where people were seen commonly eating food during pilot studies, and were undertaken at three equidistant locations along each of these streets. The Cathedral grounds, the café near St Andrews’ Harbour and the benches overlooking the Aquarium on The Scores were also locations where people were seen eating food, but because of their smaller areas, only one survey location was required at each. Surveys were undertaken on North Street, to compare with the other major streets, because this area had fewer eateries and also lower footfall. The number of people seen during each watch (recorded separately as adults and children) were recorded within a 10m radius of the watch location using clicker counters. Whether people were eating food and the type of food that they ate was noted. Attacks or attempted attacks for food were recorded, as were incidences of gulls begging for food and eating food dropped by people. The species of gull (lesser black-backed or herring gull) involved in any incidences was also recorded.

#### ii) Analysis

Rates of food theft were less common than expected so different food types were made into one variable “amount of food” and adults and children were made in to another variable “number of people”. A new dependent variable “negative human-gull interaction” was produced by combining data recorded for successful and attempted attacks for food, incidences of gulls begging for food, and gulls eating food dropped by people. Data were then analysed with GLMs (with a binomial error distribution) and models included time of day, the number of people present (adults and children), location, and amount of food. This provided information on which locations were most likely to have negative human-gull interactions, whether the number of people in an area and the amount of food they had impacted the likelihood of an interaction, and whether time of day was an important factor. Frequency of attacks were calculated by dividing the number of attacks seen by the total hours of timed watches conducted.

## 3. Results

### a) Abundance and distribution of nesting gulls

Two species of gull can be found nesting on buildings in St Andrews, herring gulls and lesser black-backed gulls. 1070 buildings across St Andrews were surveyed; a total of 46 herring gull nests and four lesser black-backed gull nests were recorded. A total of 119 herring gull chicks were seen, whilst 14 lesser black-backed chicks were recorded. Herring gulls represented 92% of the population by nests and 89.5% by chicks seen (Table 1).

**Table 1).** Nests and chicks recorded on buildings during street and vantage point surveys in St Andrews. The percentage of the overall population that each species constitutes is given as evidenced by nest numbers recorded and by presence of chicks.

| *Species* | *No. of nests* | *No. of chicks* | *% of population by nest* | *% of population by chicks* |
| --- | --- | --- | --- | --- |
| *Herring gull* | 46 | 119 | 92.0 | 89.5 |
| *Lesser black-backed gull* | 4 | 14 | 8.0 | 10.5 |

Standing gulls on buildings were also recorded throughout the street-level surveys; these were revealed to be a good indicator of the presence of nests on a building (Probability of a nest being present = 1 / (1 + exp(−((3.77*Number of standing gulls)−4.34))), z=9.0, p<0.001; Figure 1a; Model 1, Table 2). Similarly, standing gulls were shown to be a good predictor of presence of chicks on buildings (Probability of a chick being present = 1 / (1 + exp(−((4.56*Number of standing gulls)−4.31))), z=11.3, p<0.001; Figure 1b; Model 2, Table 2).

**Figure 1).**
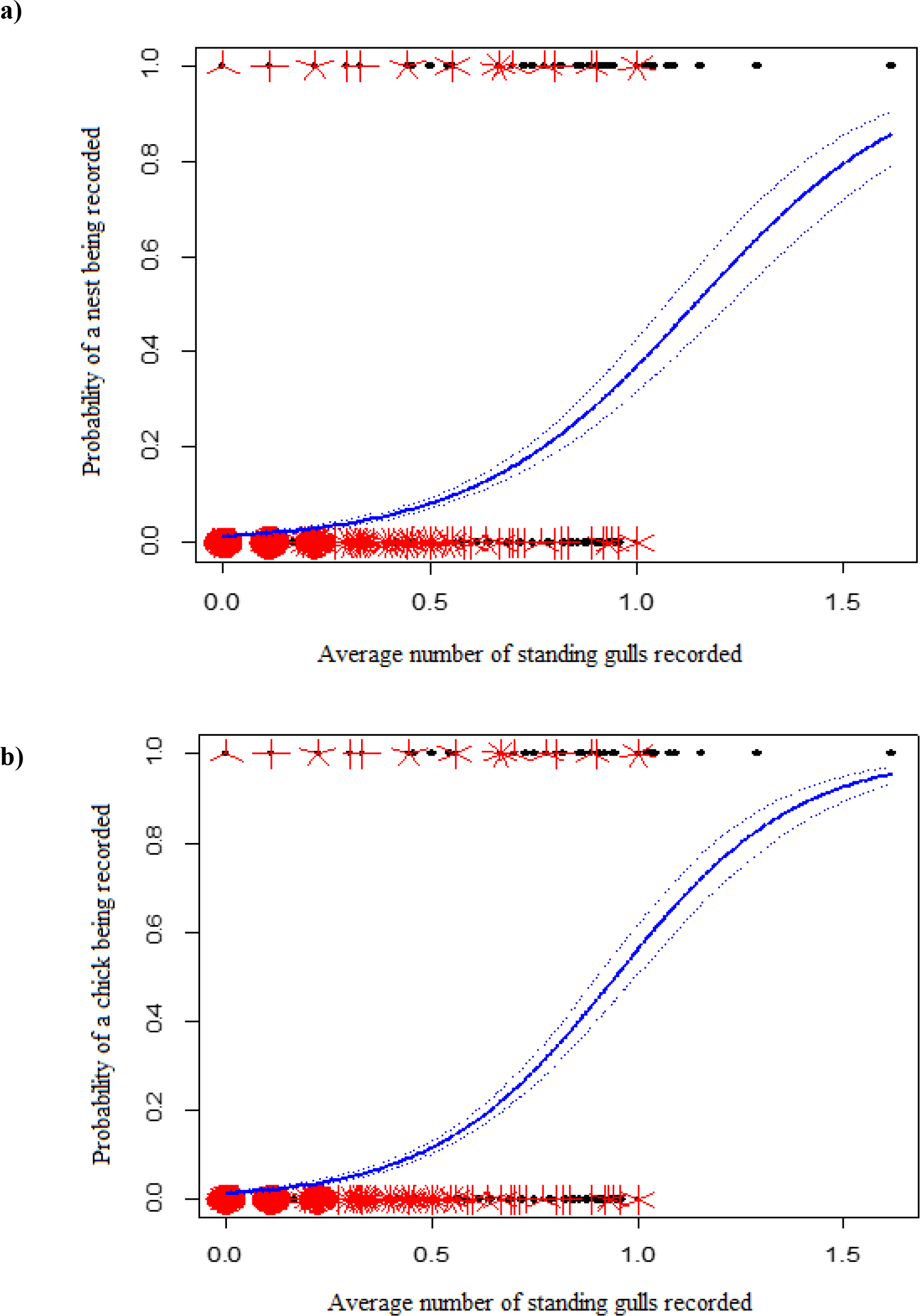
Sunflower plots showing the relationship between standing gulls and the probability of nests and chicks being recorded. Lines given ± 1SE (dotted lines); the number of petals at each point represents an observation. (a) As the average number of standing gulls recorded on a building increased the probability of a nest being recorded also increased (Model 1, Table 2).(b) A graph showing the relationship between standing gulls and the probability of chicks being recorded (Model 2, Table 2). As numbers of standing gulls on a building increased the probability of chicks being recorded also increased.

**Table 2).** Model evaluation of the most important variables when considering the likelihood of nests, chicks, attacks on waste sources and negative human-gull interactions being seen, from best models. Models are used throughout the results section (^a^= z value, ^b^= F value, ^c^= t value) *street and location are factors so for estimates see figures 2 and 6a respectively. All models assume a binomial error distribution.

| **Model no. and dependent variable** | **Model** | **Test Statistic** | **p-value** | **R<sup>2</sup>** | **Parameter Estimates** | **Standard Error** |
| --- | --- | --- | --- | --- | --- | --- |
| **1. Probability of nest being seen** | Standing gulls | 9.0<sup>a</sup> | <0.001 | 0.31 | 3.8 | 0.41 |
| **2. Probability of chick being seen** | Standing gulls | 11.3<sup>a</sup> | <0.001 | 0.39 | 4.6 | 0.40 |
| **3. Nest on buildings** | Street | 3.5<sup>b</sup> | <0.001 | 0.14 | * | * |
| **4. Nests seen from vantage points** | Intercept | 4.8<sup>c</sup> | <0.001 | 0.46 | 0.27 | 0.056 |
|  | Nests seen at street-level | 8.6<sup>c</sup> | <0.001 |  | 0.96 | 0.11 |
| **5. Rubbish attack** | Bin bags | 15.9<sup>c</sup> | <0.001 | 0.25 | 0.19 | 0.22 |
|  | Tied gull-proof | -3.6<sup>c</sup> | <0.001 |  | -0.19 | 906.9 |
|  | Untied gull-proof | -0.2<sup>c</sup> | 0.8 |  | -0.090 | 0.81 |
|  | Permanent bin | -14.7<sup>c</sup> | <0.001 |  | -14.6 | 0.32 |
| **6. Negative human-gull interaction** | People | 7.0<sup>c</sup> | 0.008 | 0.35 | -0.017 | 0.012 |
|  | Food | 103.3<sup>c</sup> | <0.001 |  | 0.36 | 0.071 |
|  | Time | 16.8<sup>c</sup> | <0.001 |  | 8.3 | 4.8 |
|  | Julian date | 2.3<sup>c</sup> | 0.1 |  | -0.011 | 0.014 |
|  | Location | 35.8<sup>c</sup> | <0.001 |  | * | * |

The probability of finding a nest on a building depended significantly on the street where buildings were located (p<0.001, F_1,49_=3.5) (Model 3, Table 2). Many nests were found along The Pends – this area included the Cathedral grounds, some of the buildings of St Leonards School and the old town wall and gates. All the buildings surveyed on The Pends had at least one nesting gull present at some point throughout the survey (Figure 2). Buildings on Butts Wynd were also locations for a number of nests, with over 60% of buildings having a nesting gull (Figure 2). Although, nests for both species were found across the survey area, more were located within the town centre where there was a higher density of buildings than in suburban areas (Figure 3a). Chicks were seen in a greater abundance than nests and were found in locations where nests had not always been recorded (Figure 3b). In addition, some nests were likely to have been unsuccessful because no chicks were recorded at some nest locations as the season progressed (Figure 3b).

**Figure 2).**
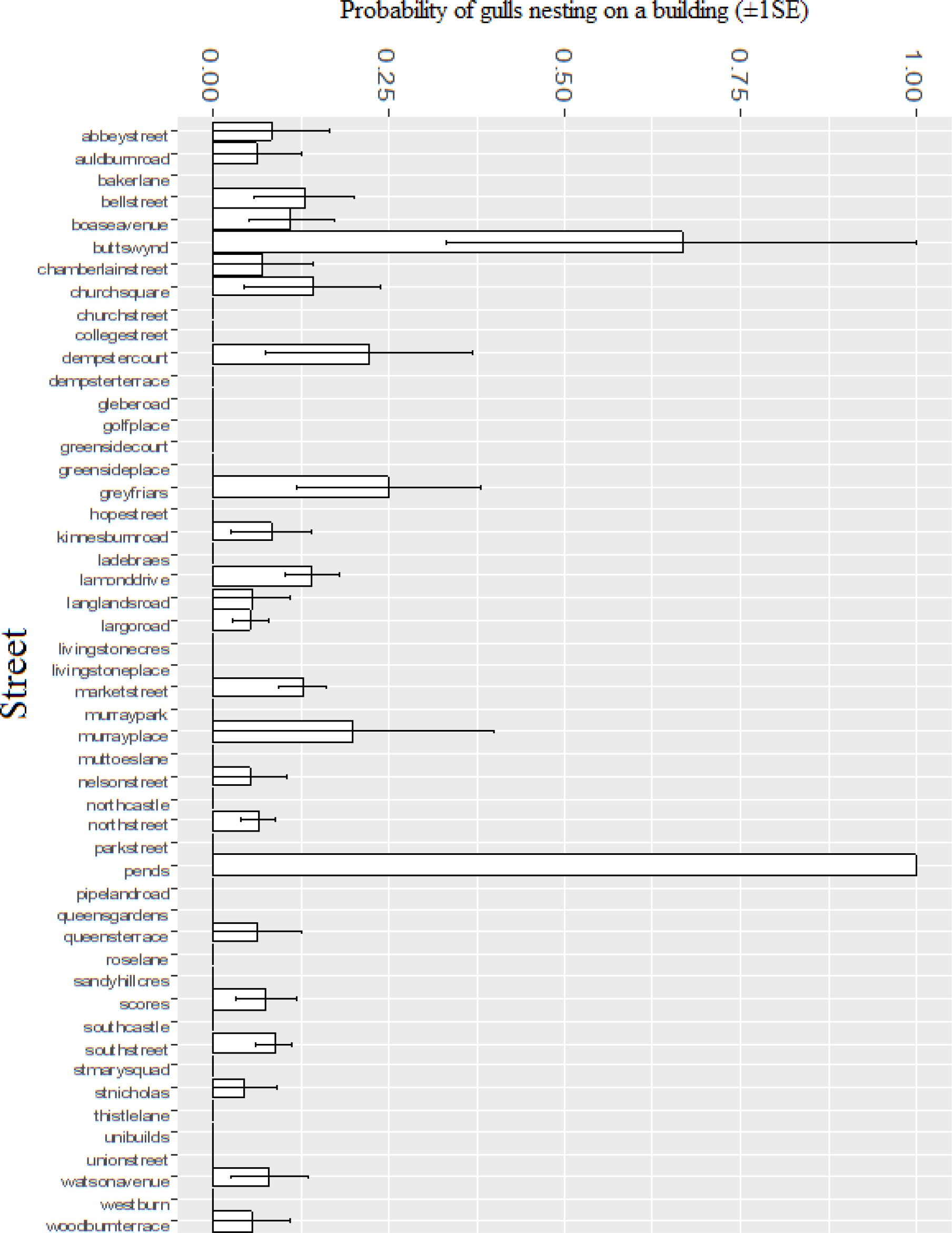
The probability of finding a nest on the different streets surveyed in St Andrews (±1 SE). Street names are given, “unibuilds” refers to the University buildings between The Scores and North Street (the Main Library, St Salvator’s Quadrangle and the Arts Building), St Mary’s quad is located between South Street and Queens Terrace. Buildings on Butts Wynd and The Pends are most likely to have nesting gulls, whilst some streets (mostly outside of the town centre) are likely to have no nesting gulls present (Model 3, Table 2).

Overall there were 93 buildings with nests and/or chicks recorded on them; 12 (13%) had a nest but no chicks (Figure 3b), suggesting that this minimum percentage of buildings lose their nests (a minimum because some nests that were never recorded will have failed). The correction factor for nests counted from vantage points relative to street estimates was 0.27, meaning that for every building checked only from the street, 0.27 nests were missed, or there was approximately one nest missed for every four buildings checked (Model 4, Table 2). This correction factor from the street-level survey/high vantage points then suggests that there were actually 64 nests ((50 from streel level *0.27) + 50). But as mentioned above, for some buildings where nests were not recorded, chicks were also recorded: overall 44 additional buildings had chicks on them where no nests were recorded. This means that a minimum of 31 nests were missed during the nest stage, assuming that chick presence on a building exactly reflected nest presence ((133 (chicks seen) / 93 (buildings with nests/chicks))*31 = 44). As already mentioned, however, some nests (an estimated 15 after applying the correction factor to the 12 nests recorded that produced no chicks) must have failed before the chick stage, so the total of nesting pairs (or nests) in the survey area was a minimum of 110 (these additional predicted nests are plotted on Figure 3b as purple triangles, with placement to reflect observed density). Therefore of the 1070 buildings surveyed in the central area of St Andrews, this study reveals that a minimum of just over 10% had nesting gulls present.

**Figure 3).**
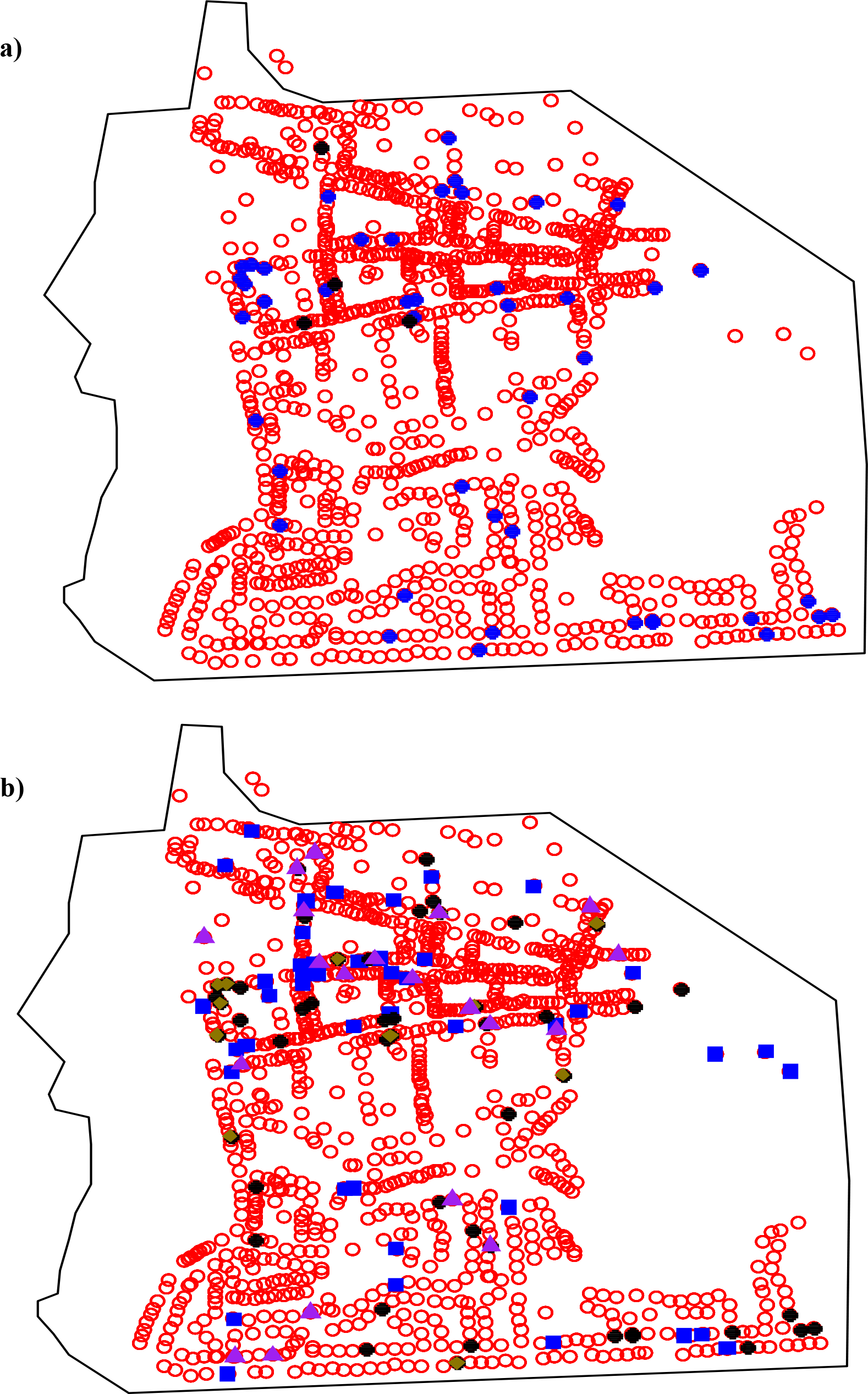
(a) A map showing nests of both herring gull (blue circles) and lesser black backed gulls (black circles) on buildings surveyed in St Andrews (red circles). (b) A map showing locations of all buildings with nests and chicks (black circles), buildings where nests were recorded but no chicks seen (gold diamonds) and buildings where only chicks were recorded but no nests had been seen (blue squares). Predicted missed nests are shown as purple triangles plotted as random points according to density of found nests.

### b) Factors determining whether rubbish sources are attacked by gulls

Gull-proof hessian bags issued by the local council were seen to be used very rarely throughout St Andrews. Of the 57 transects conducted in which bin bags, gull-proof bags and permanent bins were recorded, gull-proof bags accounted for only 12.1% of bags seen – with 87.9% of bin bags being left unprotected. Of the gull-proof bags seen only 39% were secured correctly; only 4.7% of total bags surveyed were securely tied in gull-proof bags. Of this 4.7%, no attacks on rubbish had been made. There was a significant difference in the overall likelihood of attack for different types of rubbish (F_3,2215_=77, p<0.001; Model 5, Table 2). Post-hoc Tukey’s HSD tests at the 0.5 level of significance and t-tests showed that there was a significant difference in the likelihood of attack between bin bags and tied gull-proof bags (t_130_ = 5.5, p<0.001), but not between bin bags and untied gull-proof bags (t_12_=0.1, p=0.99). There was also a significant difference between tied and untied gull-proof bags (t_10_=-1.5, p<0.05), and also between untied gull-proof bags and permanent bins (t_10_=1.4, p<0.001), other differences were not significant.

Permanent bins were seen to be accessible and attacked by gulls throughout the survey; these were shown to be less vulnerable to attack than bin bags that were not contained within gull-proof bags (t_131_=5.2, p<0.001) (Figure 4). Although attacked less often, permanent bins when attacked were noted to provide significant sources of waste on the streets – this was then cleared daily by council litter picking services. Permanent bins were present across the town centre, and both gulls and carrion crows (*Corvus corone*) were seen removing litter and food from these bins, a map of locations for permanent bins in the town centre can be found in the appendix (Figure B). Observations of attacks on commercial and private wheelie bins were also noted, with both gulls and carrion crows seen removing waste from wheelie bins where lids were left open.

**Figure 4).**
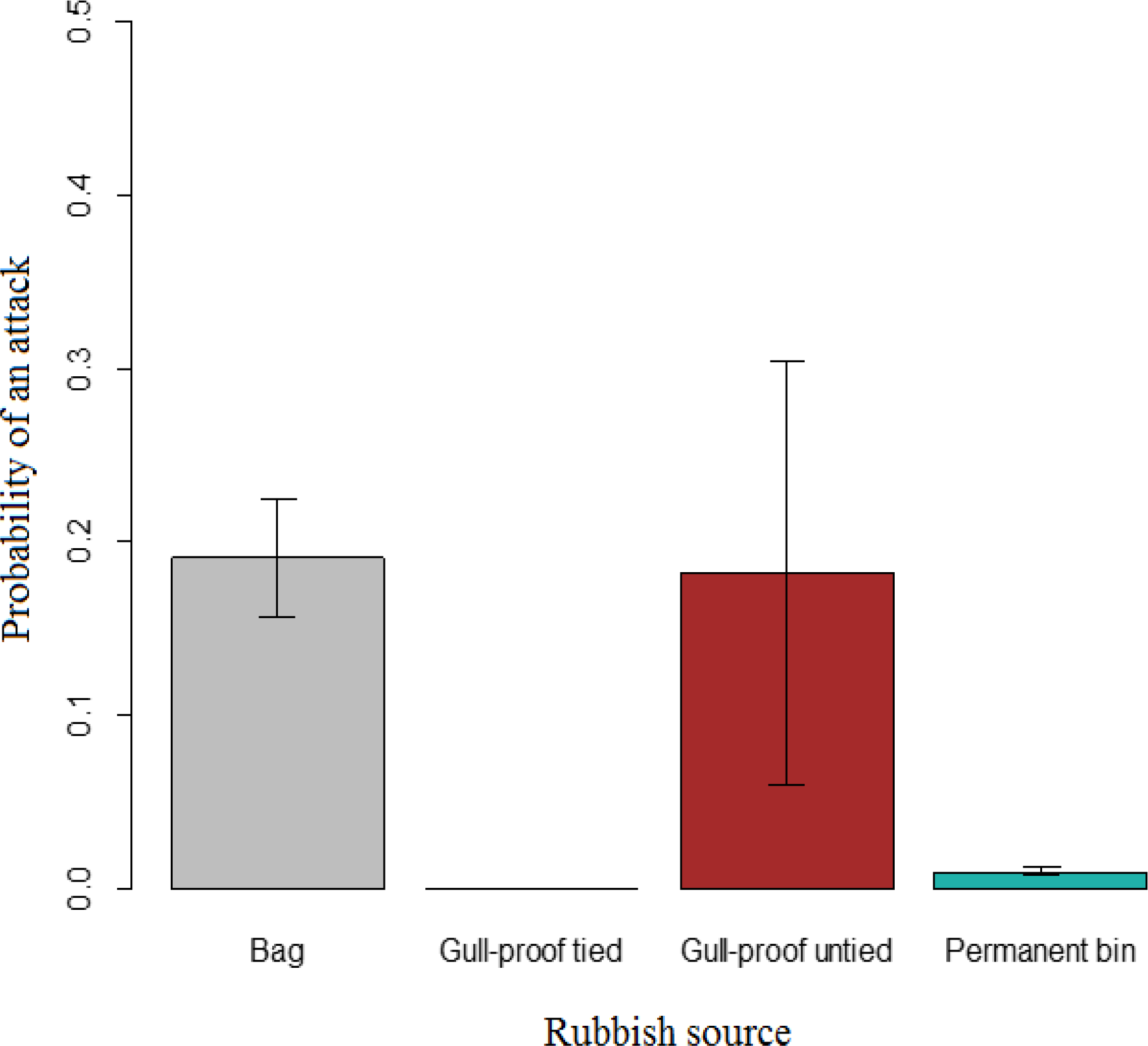
Probability a rubbish source being attacked by any gull species on transects of streets in St Andrews (error bar = ± 1SE) (Model 5, Table 2). There was a significant difference in the probability of attack between rubbish sources. Unprotected bin bags had the highest mean probability of being attacked (0,19), whilst putting a bag inside a gull-proof hessian bag was only effective when the gull-proof bag was secured properly; tied gull-proof bags had 0 probability of attack whilst untied had a similar probability (0.18) of attack to bin bags. Permanent bins had a relatively low probability of attack (0.01).

### c) The frequency and probability of negative human-gull interactions involving food

During the 10-week survey period, successful attacks were only witnessed by the surveyor on eight occasions. 11 attempted attacks in total were seen across all locations during the survey; this equates to roughly one attempt to steal food every 3.1 hours on average; whilst a successful attack was seen once every 4.25 hours on average. Location was shown to be important, with some sites seeing attempts and successful attacks more frequently, and other sites never seeing attempts or attacks (Figure 5). Location (p<0.001, F_9,970_=35.9), time of day (p<0.001, F_1,970_=16.8), and number of people present (p<0.05, F_1,970_=7.0) were shown to be significant indicators of a negative human-gull interactions being observed (Model 6, Table 2). Location 10 witnessed most attempts and this was on The Scores at the benches overlooking the aquarium – here attempts occurred 2.6 times every hour, whilst in the Cathedral grounds (location 8) an attempt occurred 0.75 times in an hour and at location 5 on Market Street, attempts were witnessed about once every 2 hours (Figure 5a). Successful attacks were seen in the same locations where attempted attacks occurred but at a lesser rate (Figure 5b). A successful attack occurred 2.3 times an hour at The Scores, once every 4 hours at the cathedral, and about once every 5 hours near on Market Street (location 5).

**Figure 5).**
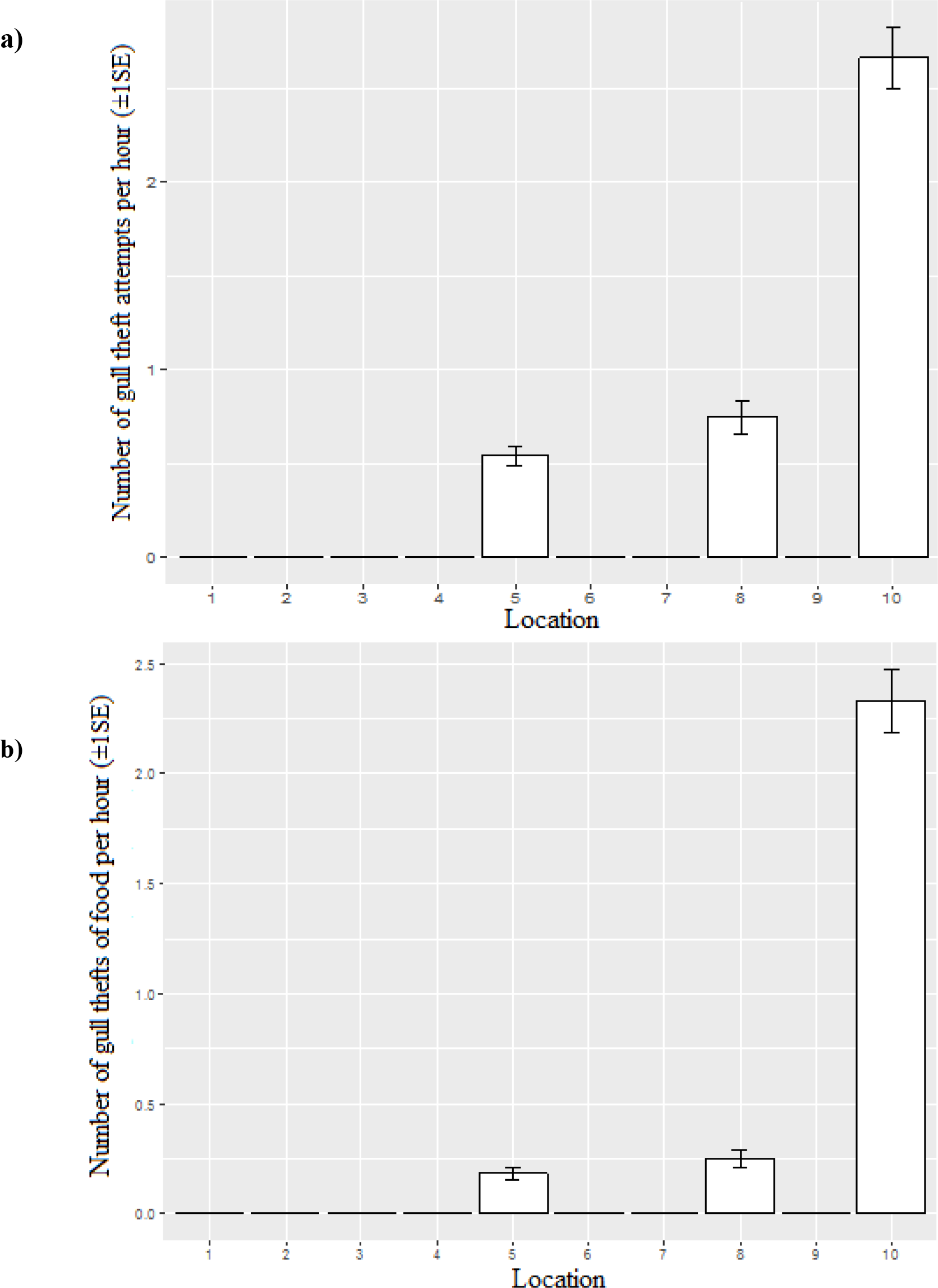
(a) The number of attempted thefts of food from people at varying locations (± 1 SE), and (b) the number of gull thefts per hour across locations in St Andrews (± 1 SE); location 5 is outside Waterstones book shop on Market St where 0.4 thefts occurred per hour; location 8 is within the cathedral grounds at benches outside the visitor centre where 0.5 thefts per hour and location 10 was on the benches overlooking the aquarium on The Scores, where more than 2 thefts were seen per hour. At other locations no gull thefts of food were seen, these included locations across North St, Market St, South St and also at the Harbour.

Probability of a negative human-gull interaction varied with food abundance, attacks were most likely where relatively low amounts of food were recorded, with between one and seven people with food (Figure 6a). However, the location of people with food was also important as only three locations (The Scores, the Cathedral grounds, and location 5 (Market St)) saw interactions. The probability of negative human-gull interactions also depended on time, with most recorded incidences falling just after midday and predictably through the lunch period, where most people would be consuming food, however SEs were large suggesting that the model result with respect to time is not robust (Model 6, Table 2; Figure 6b). The relationship between a negative human-gull interaction and the number of people (Figure 6c), shows negative interactions are most likely when between 10-50 people (a relatively low number) are in sight from a watch location.

**Figure 6).**
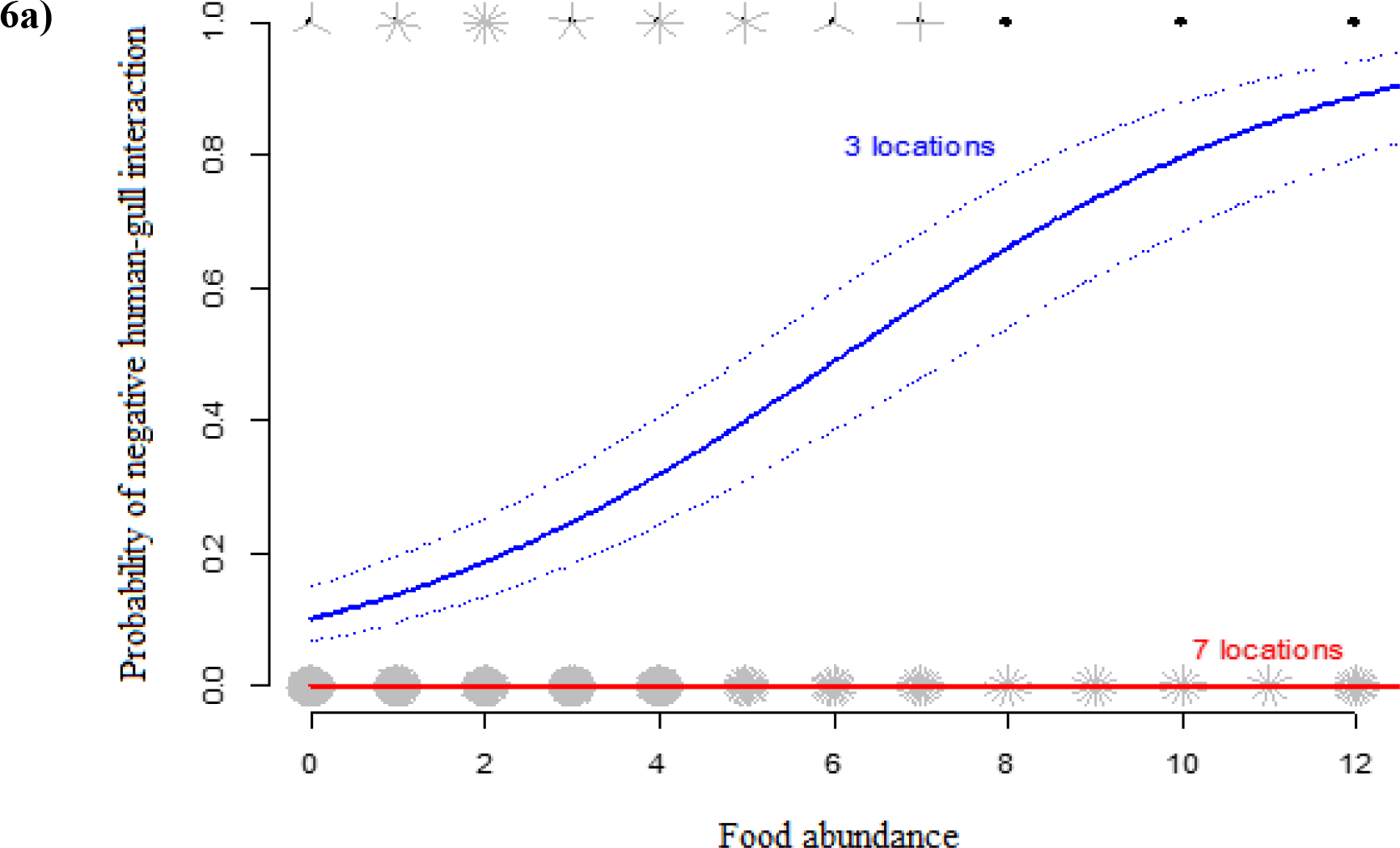
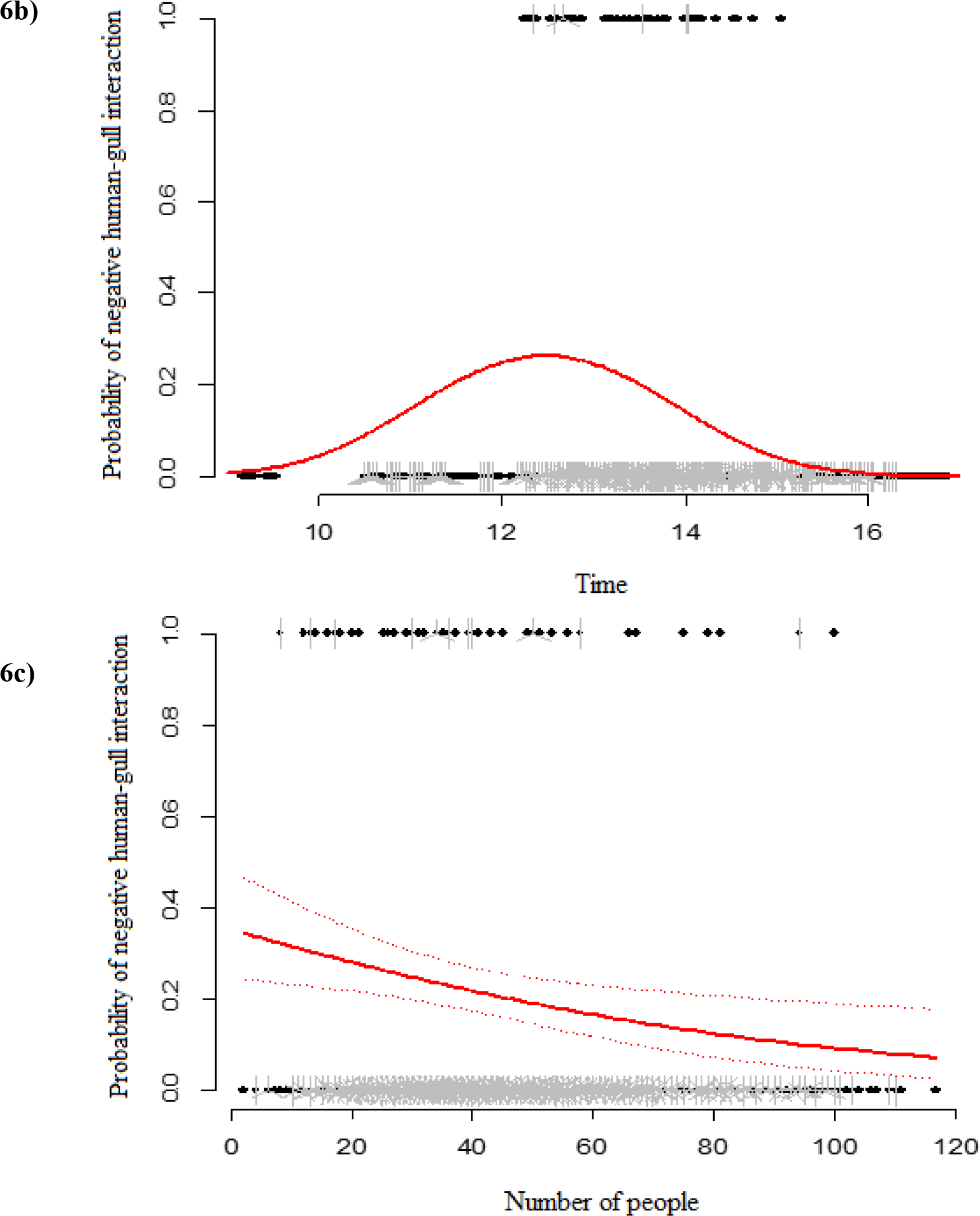
Sunflower plots showing the relationships between food abundance, time of day, and number of people present with the likelihood of a negative human-gull interaction (successful and attempted attacks on people for food, gulls begging for food, and gulls feeding on food dropped by people(Model 6, Table 2). Each petal in the plot represents an observation. All graphs are given ± 1SE (represented by dashed lines). (a) Probability of negative interaction with food abundance (the number of people with food in a 10 minute survey). The blue line shows the predicted probability of a negative interaction in the 3 locations (The Scores, The Cathedral and at a location on Market Street) where negative interactions occurred; the red line shows the attack rate at the 7 locations where no interactions were seen. (b) Probability of a negative human-gull interaction was most likely at about midday and over the lunch time period, time of day is significant in likelihood of seeing an attack. Note SEs are at 0 and 1 suggesting the model result with respect to time is not robust. (c) Probability of negative human-gull interactions is highest between 10-40 people passing through an area.

## 4. Discussion

The aim of this study was to quantify three aspects of the human-gull problem in St Andrews: the density, distribution and success of nesting gulls; the probability of different types of waste sources being used by foraging gulls; and the frequency and probability of negative human-gull interactions including gulls taking food from people. Overall, results suggest that actual problems with gulls may not be as significant as expected; human perceptions of issues in St Andrews may be disproportionate to the results of this study because of negative preconceptions already attached to these species (Dickman, 2010). A minimum of 110 gull nests were estimated across the survey area, which represents nests on only ~10% of buildings. Gulls were seen feeding opportunistically on permanent bins and unprotected bin bags. Securing waste effectively prevented gulls from accessing and spreading litter. Incidences of gulls attacking people for food were rare. However, most people seem to have a “gull story” to tell, suggesting that these events are particularly memorable when they do occur, rather than reflecting their frequency. This study shows the importance of quantifying issues and highlights that human perceptions may not align with quantified results.

### a) Nesting gulls

In St Andrews, herring gulls were seen nesting in the town in higher numbers than lesser black-backed gulls and accounted for 92% of the population by nests seen. This is similar to results from existing urban roof-nesting surveys, in which herring gulls have constituted a higher proportion of mixed urban colonies than their lesser black-backed counterparts (Cramp 1971; Raven and Coulson, 1997, Seabird Census, 2004). This is potentially because lesser black-backed gulls came to these advantageous environments later than herring gulls, and originally joined towns and cities already colonised by other gull species (Raven and Coulson, 1997). However, numbers of lesser black-backed gulls seen in towns – including St Andrews – may soon be similar to herring gulls; the rate at which these gulls are moving in to urban habitats is increasing faster than the rate for herring gulls and they have been found colonising urban areas further inland (Raven and Coulson, 1997; Nager and O’Hanlon, 2016).

Estimates of roof-nesting gulls showed that almost 90% of the buildings in St Andrews were unoccupied, suggesting that the town has the capacity to support considerably higher numbers of nesting gulls in the future. This may be an issue if lesser black-backed gulls continue increasing in numbers and moving into urban environments. More gulls were seen nesting on older buildings in the town centre than on modern buildings in the suburbs. Buildings in the town centre may be closer to natural nesting sites and food resources; buildings on Butts Wynd and The Pends had a high probability of nests being recorded. At these locations, buildings were likely to be attractive to nesting gulls as no visible gull management was in place; they are tall and resemble gulls’ natural nesting habitats on cliff ledges (JNCC, 2016a, b). In contrast, the more modern buildings in the suburbs were home to less nesting gulls, perhaps because they provided fewer stable external structures for nesting than older buildings with multiple chimney stacks and crevices for shelter. In all cases where nests were seen on buildings, they were either between chimney pots, against chimney stacks, or on flat roofs against an external wall. Previous studies have found that gulls tend to nest successfully adjacent to solid structures (Belant, 1993) and results from St Andrews support this.

Results from the data collected on nesting gulls across St Andrews’ town centre and suburbs show that just over 10% of buildings have nesting gulls on them. Understandably, many people do not want gulls on their roofs; they can be noisy, damage property and also be potentially violent towards residents particularly during the nesting season (Rock, 2005). Results from this study are important when considering management measures to prevent gulls nesting on roofs. Currently various methods exist to prevent nesting; these include the use of wires, nets and mylar flags (Blockpoel andTessier, 1984; Belant and Ickes, 1996, 1997), some of these methods were observed on buildings in St Andrews (the University Main Library uses nets to prevent nesting and loafing by gulls). Other residents have resorted to make-shift methods, for example one house on Lamond Drive has barbed wire wrapped around chimney pots to deter gulls from nesting. However, this was not effective as a herring gull was seen nesting between the barbed wire and successfully hatched two chicks. Nevertheless, because of the number of unoccupied roofs available in St Andrews, management to move nesting gulls from one roof would simply shift the issue to a nearby building. Management of roof-nesting in St Andrews is unlikely to be successful unless applied to all buildings, which would be highly costly. Any new developments of buildings should consider planning to make them less attractive to gulls, using dark roofing materials and avoiding the provision of stable structures for nesting where possible (Belant and Dolbeer, 1993; Blokpoel and Tessier, 1991). Lethal management is not an option for these species as they are protected and we have a conservation responsibility for them. Gulls are long-lived, and once they have colonised a town – as they have in St Andrews – it is likely they will continue to breed there throughout their lives (Chabrzyk and Coulson, 1976; Bosman et al., 2016).

Comprehensive and cost-effective methods for estimating urban gull numbers are still in development and this survey may have underestimated gull numbers, because even with vantage surveys and an applied correction factor it is likely some nests will be unaccounted for. A study by Coulson and Coulson (2015) assessed effectiveness of correction factors compared to using a cherry-picker to get accurate counts of nests on each building. They found that correction factors underestimated numbers, showing approximately 84% of nests in their particular survey area; this is likely to vary between locations because different sites have different attributes and visibility. Using a cherry-picker is likely to be very costly, and unfeasible in most surveys. Conducting street and vantage point surveys as in this study can be time-consuming, hence incidences of standing gulls were recorded to see whether they were a good indicator that a nest would be present on a building. A relationship was seen between the two, but this is unlikely to accurately estimate nesting gulls. Nevertheless it could provide a quick way to estimate minimum nest numbers in pilot studies and assess the need for more comprehensive surveys. Even if overall nest numbers were underestimated in this survey, it would not change that the majority of buildings do not have gulls nesting on them; the same issues with management would remain.

### b) Vulnerability of rubbish sources to attack

Success of nesting gulls has been attributed to additional food availability in urban environments from anthropogenic sources (Rock 2005, Monaghan, 1979), and gulls have been shown to exploit individualistic foraging behaviours in a recent GPS tracking study (Rock et al., 2016). Transects conducted throughout the survey period showed that black bin bags had the highest probability of attack. Very few hessian gull-proof bags issued by the local council were seen to be used and secured properly throughout the study, but where they were secured, none of these were attacked. In contrast, where these bags were not tied correctly they were almost as likely to be attacked as completely unprotected bin bags. The probability of attack on permanent bins was lower than that seen for bin bags and unsecured gull-proof bags. This may be because bins were attacked less frequently, but it may be that these rubbish sources were more frequently emptied than others or were slightly more difficult to access for large gulls. Furthermore, council employees were frequently seen clearing up rubbish that had been removed from these permanent bins by gulls and carrion crows. Although, probability of attack was less, spread of waste from these sources was high – preventing gulls accessing this rubbish would also prevent other urban species such as carrion crows from foraging on this waste. Discerning whether waste sources were attacked by gulls or other foraging urban wildlife such as carrion crows was difficult, but on the majority of occasions that rubbish was disturbed, damage was consistent with a gull attack. Nevertheless, problems with litter spreading are independent of the species attacking the waste, and securing rubbish is likely to be beneficial against all foraging animals. Interestingly, carrion crows and herring gulls were observed together in groups accessing food from bins; on more than one occasion smaller crows were seen to remove food from bins, before gulls took the food. This may be an example of gulls’ cognitive abilities indicating their capacity to learn how to most efficiently access anthropogenic food.

This conflict can be managed in several ways. Firstly, results show that placing bin bags in hessian gull-proof bags is an effective way to prevent the spread of litter; encouraging individuals to do so should help to resolve this problem. However, these bags must be secured properly, as they have little effect, if any, when left untied. Increased collection of rubbish from permanent bins will also help to mitigate the problem. Although these bins were seen to be attacked less frequently, when they became full it was noticed that both gulls and carrion crows could access them and spread litter. Permanent bins that are closed off to gulls by a flap or another method would also help mitigate the problem of gulls foraging on these sources and prevent these bins providing a constantly available food source to gulls in St Andrews.

### c) Human-gull conflict involving food at street-level

Despite numerous reports of frequent thefts of food, this study revealed this to be a very rare event. In total, attempted attacks on people for food by gulls were only seen 11 times in 30 hours of timed watches over a 10-week period. These incidences were only seen at three of the 10 monitored locations. Negative interactions with gulls (including attacks for food) were most likely at The Scores over the lunch time period; it may be that some gulls have learnt that this is a good location to target individuals for food. The number of people at a site was seen to influence the likelihood of interactions between people and gulls, with locations with relatively low numbers of people vulnerable to incidences. In addition, at The Scores and in the Cathedral grounds where attacks occurred, people were seen feeding gulls. Gulls were also observed begging for food, potentially an adjustment of natural begging behaviour to adults for food turned to people; an example of gulls’ plastic foraging behaviours (Morand-Ferron et al., 2007). The third location where an incident occurred was on Market St near the Waterstones book shop. This was close to a café with high footfall where gulls may have learnt that leftovers and food from people are readily available. Time of day is likely to have some impact on when gull attacks occur, but this is probably linked to periods of time when people are eating on the street – such as lunch time.

Management of this issue is difficult because these events are rare, but perceived to be regular occurrences because incidences are memorable. Nevertheless, some management may be possible, particularly in areas where higher rates of theft and negative interactions were seen. These could involve educational signs about gulls and their conservations statuses, but warn individuals to be vigilant when eating and not to feed gulls in these areas. If food sources are removed or made more difficult to access it is likely that gulls will look elsewhere for food and cause less harassment to people.

### d) Conclusions and Recommendations

The aim of this study was to quantify three major aspects of the human-gull conflict in St Andrews. By quantifying these, a number of management practices to reduce human-gull conflicts can be suggested: where bin bags are used they should be secured in hessian gull-proof bags provided by the council and tied up properly to avoid spread of litter. A greater number of permanent bins would help to reduce the problem of overfilling and limit the availability of waste. Furthermore, permanent bins with flaps or enclosed disposal systems, for example, “Bigbelly bins” used in the local town of Anstruther (Bigbelly, 2016) would prevent gulls and other foraging animals accessing these bins; a bin of this type is currently being trialled on Market St (D Angus 2017, personal communication, 25 January). Although few incidences of attacks on people for food were seen, some locations were shown to be more prone to incidences than others. Signs providing warnings about gulls taking food at these sites and discouraging people from feeding gulls in these areas may help to mitigate problems. Outside of the breeding season, foraging by gulls on anthropogenic food sources may be different and further investigation into human-gull interactions during this period are necessary.

It is likely that the key issue to solving the St Andrews “gull problem” is to break the culture of having anthropogenic food sources available to gulls. This requires a multi-faceted approach, applying the recommendations of this survey to make access to waste and food more difficult for gulls in St Andrews. There is unlikely to be a solution to remove or reduce numbers of nesting gulls as there are so many vacant available sites for nests. Management may be successful for individual buildings, but will only shift the problem to an adjacent or nearby building. Management over the whole of St Andrews would be unfeasible because of high costs and would probably shift the colony to another nearby town and lethal control is not an option because these animals have protected status and are of conversation concern. Increases in roof-nesting gulls may be slowed by altering human behaviours to reduce availability of food resources. However, St Andrews is at a coastal-urban interface and thus food is still available from natural environments, so reducing anthropogenic food may only have limited success.

The gull problem in St Andrews may not be as significant as first thought; it appears that people have misconceptions surrounding the presence and behaviour of gulls in the town and the frequency of events. Altering human perceptions can be difficult, but teaching the public more about the two gull species found in St Andrews and their protected status may help. The name “seagull” is commonly used as a term for all gulls, but seeing them as individual species (herring gulls and lesser black-backed gulls) may help to forge more positive human-gull interactions. It has been highlighted in the past that personal connections with animals and greater knowledge of their biology and potential threats they face can alter individuals’ views (Jerolmack, 2008; Dickman, 2010). Where species are of conservation concern and in conflict with humans, education by enthusiastic and passionate individuals may be the best way to change peoples’ perceptions (Bonta, 2008; Dickman, 2010).

## 6. Acknowledgements

I thank Professor Will Cresswell, the St Andrews Business Improvement District (BID), organisations and individuals across St Andrews for providing me with access to vantage points for surveys, members of the University’s Estates Team, particularly Kevin Litster, St Leonards School and Simon Foxx, the team at Holy Trinity Church, Historic Environment Scotland, and Naomi Boon.

## 7. Appendix

**Figure A).**
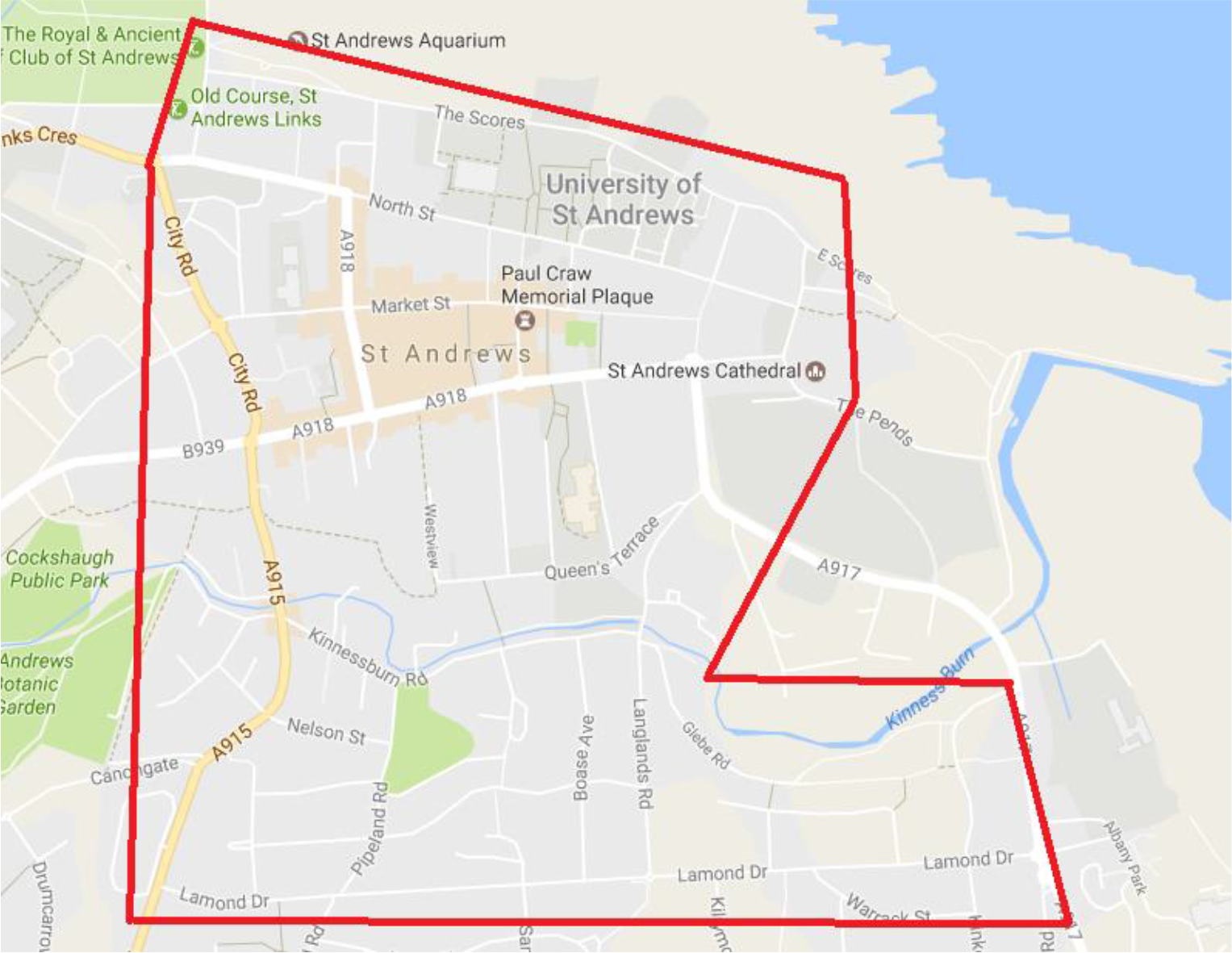
A map of the survey area showing streets that were surveyed during data collection for nesting gulls. Map adapted from Google Maps (2017).

**Figure B).**
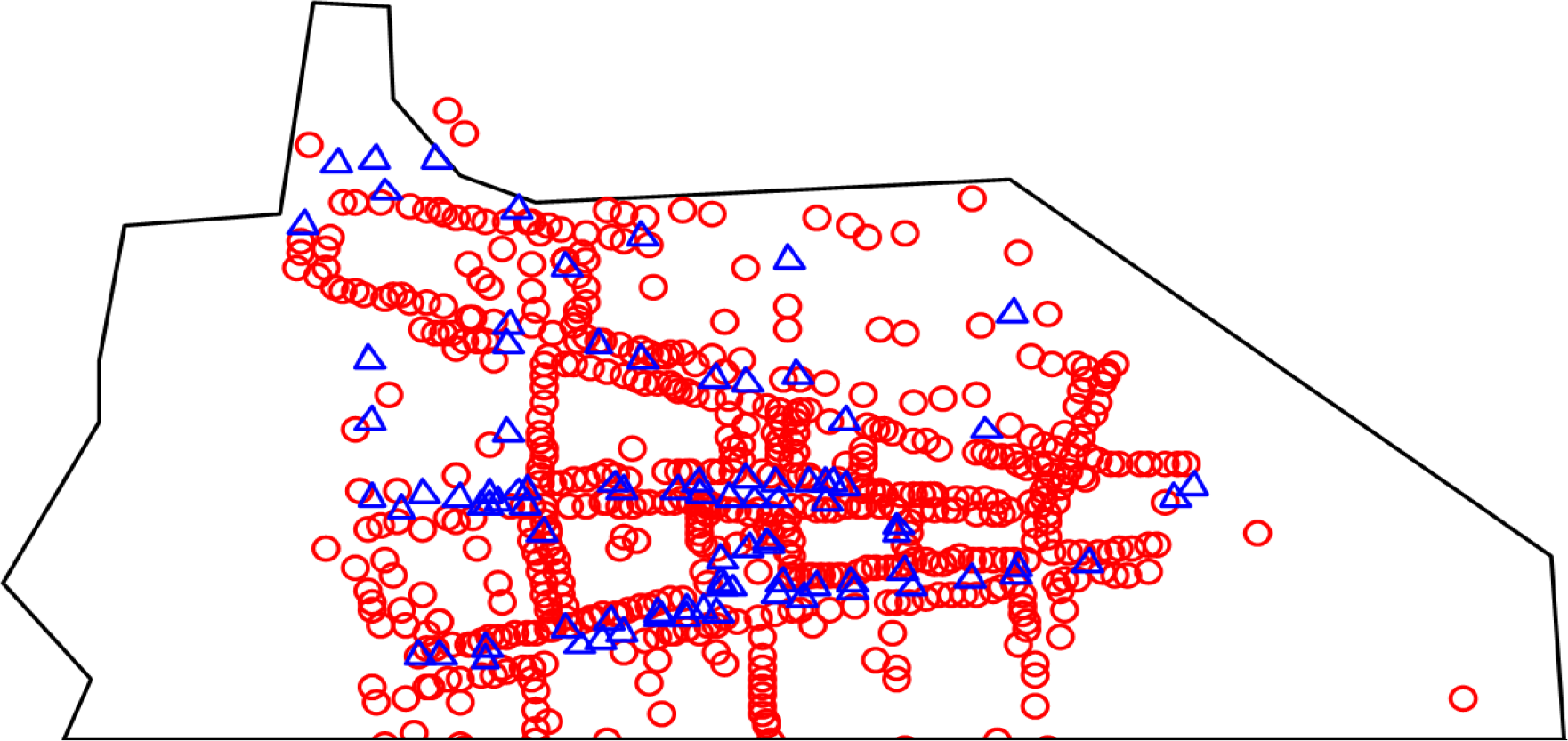
A map showing the locations of all permanent bins (blue triangles) in St Andrews town centre in relation to buildings (red circles). 75 permanents bins are located in the town centre.

**Table A).** A table giving details on locations (including coordinates) for sites where timed watches were conducted.

| *Location Number* | *Street/area* | *Coordinates* |
| --- | --- | --- |
| 1 | South Street | 56.33964053, -2.79175758 |
| 2 | South Street | 56.33946213, -2.79499769 |
| 3 | South Street | 56.33899234, -2.79872061 |
| 4 | Market Street | 56.34032885, -2.79831827 |
| 5 | Market Street | 56.34043440, -2.79668212 |
| 6 | Market Street | 56.34032142, -2.79582918 |
| 7 | North Street | 56.34137840, -2.79559314 |
| 8 | Cathedral | 56.33983826, -2.78876424 |
| 9 | Harbour | 56.33893287, -2.78366804 |
| 10 | Scores | 56.34331539, -2.80040503 |

## References

Abay GY, Bauer H, Gebrehiwot K and Deckers J. (2011). Peri-urban spotted hyena (*Crocuta crocuta*) in northern Ethiopia: diet, abundance and economic impact. European Journal of Wildlife Research 57:759–765.

Adams LW. (2005). Urban wildlife ecology and conservation: a brief history of the discipline. Urban Ecosystems 8:139–156.

Atkins A, Redpath SM, Little RM. and Amar A. (2017). Experimentally manipulating the landscape of fear to manage problem animals. The Journal of Wildlife Management 22:1–7.

Belant JL. (1993). Nest-site selection and reproductive biology of roof- and island-nesting herring gulls. Transactions of the North American Wildlife and Natural Resources Conference 58:78–86.

Belant JL. (1997). Gulls in urban environments: landscape-level management to reduce conflict. Landscape and urban planning 38(3-4):245–58.

Belant JL and Dolbeer RA. (1993). Population status of nesting laughing gulls in the United States 1977-1991. American Birds 47(2):220–4.

Belant JL and Ickes SK. (1996). Overhead wires reduce roof-nesting by ring-billed gulls and herring gulls. Proceedings of the Vertebrate Pest Conference 17:112–116.

Belant JL and Ickes SK. (1997). Mylar flags as gull deterrents. Great Plains Wildlife Damage Control Workshop Proceedings 359:73–80.

Belant JL, Ickes SK and Seamans TW. (1998). Importance of landfills to urban-nesting herring and ring-billed gulls. Landscape and urban planning 43(1):11–9.

Bigbelly. (2016). Waste & Recycling Stations, A Solution for Every Corner. Available from: http://bigbelly.com/solutions/stations/ [Accessed 21 February 2017].

Blokpoel H and Tessier GD. (1984). Overhead wires and monofilament lines exclude ring-billed gulls from public places. Wildlife Society Bulletin 12(1):55–8.

Blokpoel H and Tessier GD. (1991) Control of ring-billed gulls and herring gulls nesting at urban and industrial sites in Ontario, 1987-1990. Fifth Eastern Wildlife Damage Control Conference 5:51–57.

Bomford M and O’Brien PH. (1990). Sonic deterrents in animal damage control: a review of device tests and effectiveness. Wildlife Society Bulletin 18(4):411–22.

Bonta M. (2008). Valorizing the relationships between people and birds: Experiences and lessons from Honduras. Ornitologia Neotropical 19:595–604.

Bosman DS, Stienen EW and Lens L. (2016). Sex, growth rate, rank order after brood reduction, and hatching date affect fìrst-year survival of long-lived Herring Gulls. Journal of Field Ornithology 87(4):391–403.

Calladine JR, Park KJ, Thompson K and Wernham CV. (2006). Review of urban gulls and their management in Scotland. A report to the Scottish Executive. Edinburgh.

Campbell MO. (2007). An animal geography of avian ecology in Glasgow. Applied Geography 27(2):78–88.

Chabrzyk G and Coulson JC. (1976). Survival and recruitment in the Herring Gull *Larus argentatus*. The Journal of Animal Ecology 45(1):187–203.

Conover, MR. (2001). Resolving human–wildlife conflicts: the science of wildlife damage management. First Edition. Florida: CRC Press. pp 1–16.

Coulson JC and Coulson BA. (2008). Lesser Black-backed Gulls *Larus fuscus* nesting in an inland urban colony: the importance of earthworms (Lumbricidae) in their diet. Bird Study 55:297–303.

Coulson JC and Coulson BA. (2015). The accuracy of urban nesting gull censuses. Bird Study 62(2):170–6.

Cramp S. (1971). Gulls nesting on buildings in Britain and Ireland. British birds 64:476–487.

Dickman AJ. (2010). Complexities of conflict: the importance of considering social factors for effectively resolving human-wildlife conflict. Animal conservation 13(5):458–66.

Eaton M, Aebischer N, Brown A, Hearn R, Lock L, Musgrove A, Noble D, Stroud D and Gregory R. (2015). Birds of Conservation Concern 4: the population status of birds in the UK, Channel Islands and Isle of Man. British Birds 108:708–46.

Environmental Systems Research Institute (ESRI). (2014). ArcGIS Release 10.3. Redlands, CA.

Erickson WA, Marsh RE and Salmon TP. (1990). A review of falconry as a bird-hazing technique. Proceedings of the Vertebrate Pest Conference 14:314–316.

Found R and Boyce MS. (2011). Predicting deer–vehicle collisions in an urban area. Journal of Environmental Management 92(10):2486–93.

Google Earth. (2016). Google Earth 7.1, St Andrews, Scotland, 56°20’27.71”N, 2°47’53.20”20”W, elevation 27m. Available from: https://www.google.com/earth/index.html. [Accessed 20 May 2016].

Google Maps. (2017). Map of St Andrews [online]. Google. Available from: https://www.google.co.uk/maps/place/St+Andrews/@56.3365279,-2.8427956,13z/data=!3m1!4b1!4m5!3m4!1s0x488650981912f9fb:0x5526f99ec05ca9f0!8m2!3d56.3397753!4d-2.7967214. [Accessed 24 January 2017].

Gusset M, Swarner MJ, Mponwane L, Keletile K and McNutt JW. (2009). Human–wildlife conflict in northern Botswana: livestock predation by endangered African wild dog Lycaon pictus and other carnivores. Oryx 43(1):67–72.

Jerolmack C. (2008). How pigeons became rats: The cultural-spatial logic of problem animals. Social problems 55(1):72–94.

Joint Nature Conservation Committee (JNCC). (2009). Directive 2009/147/EC on the conservation of wild birds. Available from: http://jncc.defra.gov.uk/page-1373. [Accessed 18 February 2017].

Joint Nature Conservation Committee (JNCC). (2016a). Herring Gull Status and Trends. Available from: http://jncc.defra.gov.uk/page-2887. [Accessed 18 February 2017].

Joint Nature Conservation Committee (JNCC). (2016b). Lesser Black-backed Gull Status and Trends. Available from: http://jncc.defra.gov.uk/page-2886. [Accessed 18 February 2017].

Lowry H, Lill A and Wong B. (2013). Behavioural responses of wildlife to urban environments. Biological reviews 88(3):537–549.

Magle SB, Hunt VM, Vernon M and Crooks KR. (2012). Urban wildlife research: past, present, and future. Biological Conservation 155:23–32.

Michalski F, Boulhosa RL, Faria A and Peres CA. (2006). Human-wildlife conflicts in a fragmented Amazonian forest landscape: determinants of large felid depredation on livestock. Animal Conservation 9(2):179–88.

Mitchell PI, Newton SF, Ratcliffe N and Dunn TE. (2004). Seabird populations of Britain and Ireland: results of the Seabird 2000 census (1998-2002). London: T and AD Poyser.

Monaghan P. (1979). Aspects of the breeding biology of Herring Gulls *Larus argentatus* in urban colonies. Ibis 121(4):475–81.

Monaghan P and Coulson JC. (1977). Status of large gulls nesting on buildings. Bird study 24(2):89–104.

Morand-Ferron J, Sol D and Lefebvre L. (2007). Food stealing in birds: brain or brawn? Animal Behaviour 74(6):1725–1734.

Nager RG and O’Hanlon NJ. (2016). Changing numbers of three gull species in the British Isles. Waterbirds 39(sp1):15–28.

Perrins CM. (1970). The timing of birds’ breeding seasons. Ibis 112(2):242–255.

R Core Team. (2016). R: A language and environment for statistical computing. R Foundation for Statistical Computing, Vienna, Austria.

Raven SJ and Coulson JC. (1997). The distribution and abundance of Larus gulls nesting on buildings in Britain and Ireland. Bird Study 44(1):13–34.

Rock P. (2005). Urban gulls: problems and solutions. British birds 98:338–55.

Rock P. (2012). Urban gulls. Why current control methods always fail. Rivista Italiana di Ornitologia 82(1-2):58–65.

Rock P and Vaughan IP. (2013). Long-term estimates of adult survival rates of urban Herring Gulls *Larus argentatus* and Lesser Black-backed Gulls *Larus fuscus*. Ringing & Migration 28:21–29.

Rock P, Camphuysen CJ, Shamoun-Baranes J, Ross-Smith VH and Vaughan IP. (2016). Results from the first GPS tracking of roof-nesting Herring Gulls *Larus argentatus* in the UK. Ringing & Migration. 31(1):47–62.

Soldatini C, Albores-Barajas YV, Torricelli P and Mainardi D. (2008). Testing the efficacy of deterring systems in two gull species. Applied Animal Behaviour Science 110(3):330–40.

Soulsbury CD and White PC. (2016). Human-wildlife interactions in urban areas: a review of conflicts, benefits and opportunities. Wildlife Research 42(7):541–53.

Vermeer K, Power D and Smith GJ. (1988). Habitat selection and nesting biology of roof-nesting Glaucous-winged Gulls. Colonial waterbirds 11(2):189–201.

Washburn BE, Bernhardt GE, Kutschbach-Brohl L, Chipman RB and Francoeur LC. (2013). Foraging Ecology of Four Gull Species at a Coastal-Urban Interface. The Condor 115(1):67–76.

Young JC, Thompson D, Moore P, MacGugan A, Watt A and Redpath SM. (2016). A conflict management tool for conservation agencies. Journal of Applied Ecology 53(3):705–11.

